# Separation of function mutations in microcephaly protein CPAP/CENPJ, reveal its role in regulating centriole number and length

**DOI:** 10.1101/2023.02.07.527589

**Authors:** Sonal Jaiswal, Srishti Sanghi, Priyanka Singh

## Abstract

Centriole are microtubule-based cylindrical structures characterized by their definite size, and stable, slow growing microtubules. The centriole core protein CPAP/CENPJ is known to act as a molecular cap regulating centriole length by interacting with microtubule/tubulin *via* the conserved microtubule destabilizing, PN2-3, and microtubule stabilizing, A5N, domains. The C-terminus of CPAP has a conserved glycine-rich G-box/TCP domain (1050-1338 amino acids). This region is involved in centriole cartwheel assembly by interacting with the cartwheel protein STIL. However, previously reported primary microcephaly mutation mapped in the G-box of CPAP, i.e., E1235V (MCPH6) affects centriole length *via* an unknown mechanism. Recently, another primary microcephaly mutation has been mapped to this region of CPAP, i.e., D1196N. However, the effect of D1196N on CPAP functioning is not known. We simultaneously characterized these two MCPH mutations in the G-box of CPAP. We identified that despite affecting the same domain of CPAP, they affect distinct CPAP functions at the centriole. The E1235V mutation caused an overly long centriole, and the D1196N mutation increased the centriole number. Interestingly, both these mutations affect CPAP direct interaction with the cartwheel protein STIL, which is involved in CPAP recruitment to the centriole. Accordingly, the CPAP E1235V centriole localization is significantly affected at the centriole. However, CPAP D1196N can still localize to centriole at levels comparable to the wild-type CPAP. We show that CPAP utilizes an alternate CEP152-dependent route for centriole recruitment. Importantly, our work highlights the importance of the CPAP region outside direct microtubule/tubulin interacting domains in influencing CPAP activity in cartwheel assembly and centriole length. Perhaps, this is why deleterious naturally occurring missense mutations are frequently occurring in this particular region of CPAP in primary microcephaly.

## INTRODUCTION

Centrosomes are major microtubule organizing centers in many dividing animal cells. They also generate basal bodies, which extend to form axoneme of cilia and flagella. Therefore, centrosomes are required for cellular processes like division, motility, and signaling (Vertii *et al*, 2016). It is a membrane-less cell organelle, mainly consisting of a pair of microtubule based cylindrical structure at the core called the centrioles, embedded in a proteinaceous matrix referred as pericentriolar material (PCM). The mature mammalian centriole is roughly 250 nm in diameter and 450 nm in length. They exhibit a characteristic 9-fold radially symmetric arrangement of microtubule triplets (Vorobjev & Chentsov, 1982; Anderson & Brenner, 1971; Gupta & Kitagawa, 2018).

Certain evolutionary conserved proteins first identified in *Caenorhabditis elegans* and *Drosophila melanogaster* tightly regulate centriole duplication during the S-phase of the cell cycle (Pelletier *et al*, 2006; Kemp *et al*, 2004; Leidel & Gönczy, 2003; Leidel *et al*, 2005; Delattre *et al*, 2004). In human cells, the new (procentriole) centriole generation involves the proximally localized parent centriole proteins, CEP152 (functional orthologue of Asterless in flies) and CEP192 (functional orthologue of SPD-2 in worms) (Sonnen *et al*, 2013). They initiate recruitment of centrosome-specific kinase, PLK4 (functional orthologue of ZYG1 in worms and SAK in flies) adjacent to the parent centrioles (O’Connell *et al*, 2001; Delattre *et al*, 2006; Bettencourt-Dias *et al*, 2005; Habedanck *et al*, 2005). Once recruited, PLK4 brings STIL (SCL/TAL1 interrupting locus; SIL) protein at the centriole, whose functional orthologue is SAS-5 in worms and ANA-2 in flies. PLK4-mediated phosphorylation of STIL in its STAN domain at the C-terminus promotes the recruitment of central cartwheel protein SAS-6 (Dzhindzhev *et al*, 2014). The oligomerization of SAS-6 homodimers dictates the nine-fold symmetry of microtubules at the centriole (Nakazawa *et al*, 2007; Kitagawa *et al*, 2011b). The N-terminus of STIL protein contain a proline-rich conserved region (CR2) that interacts with the CPAP (centrosomal P4.1-associated protein or CENPJ) protein (Cottee *et al*, 2013). SAS4, the worm orthologue of CPAP, is shown to regulate centrosome size by regulating the levels of PCM organized around the centrioles (Kirkham *et al*, 2003). The overexpression of CPAP increases centriole length in cycling human cell lines, which translates to abnormal spindle organization and cell division (Kohlmaier *et al*, 2009; Schmidt *et al*, 2009; Tang *et al*, 2009).

CPAP was first identified in a yeast two-hybrid screen as an interacting partner of γ-tubulin complex (Hung et al, 2000). Three major conserved domains have been identified in CPAP protein, which includes the N-terminal microtubule destabilizing, PN2-3 domain (amino acid 311-422) (Hung *et al*, 2004; Cormier *et al*, 2009), followed by the microtubule-stabilizing, A5N domain (amino acid 423-607) (Hsu *et al*, 2008) and the C-terminal located T-complex protein 10 (TCP) or glycine-rich G-box domain (amino acid 1150-1338) (Zheng *et al*, 2014). Structural work revealed that the CPAP microtubule destabilizing/stabilizing domains act as a molecular cap for defining organelle size by engaging the PN2-3 domain with the β-tubulin site involved in longitudinal tubulin-tubulin interaction. It also engages the A5N domain for microtubule stability. Therefore, CPAP ensures slow and progressive growth of centriolar microtubules (Sharma *et al*, 2016). The C-terminus of CPAP (895-1338), which include the G-box domain, is involved in direct interaction with the STIL protein (Tang *et al*, 2011). CPAP protein also exhibits five short coiled-coil (CC) regions, and the CC5 region close to the G-box domain is responsible for the protein homo-dimerization (Zhao *et al*, 2010).

Abnormalities in centrosomes have been associated with human diseases like cancers, ciliopathies, and neurodevelopmental disorders. Autosomal recessive primary microcephaly (MCPH) is a neurodevelopmental disorder causing small head size and intellectual disability (Thornton & Woods, 2009). It is a genetically heterogeneous disease. However, mutations in the core centriole proteins, CEP152, PLK4, STIL, SAS-6 and CPAP have been associated with MCPH, suggesting the importance of proper centrosome organization in brain development (Jaiswal & Singh, 2021). The missense mutation, E1235V, in the G box domain of CPAP was identified in primary microcephaly (MCPH6) patients (Bond *et al*, 2005). This mutation reduces the direct interaction of CPAP with the cartwheel protein STIL (Tang *et al*, 2009). CPAP E1235V overexpression in cells has no major effect on centriole numbers, but it causes excessively long centrioles as compared to the wild-type protein. Nevertheless, such centriole defects result in defective spindle organization and positioning, possibly leading to primary microcephaly by affecting the division of neural progenitor cells in the neonatal brain (Kitagawa *et al*, 2011a). Structural work of the CPAP G-box domain from the *Danio rerio* shows its fibrillary structure with a periodicity of ~8 nM, a length comparable to the size of tubulin heterodimer and half the size of successive cartwheel spoke in *Tryconympha* basal body (Guichard *et al*, 2012; Hatzopoulos *et al*, 2013). This suggests that CPAP is not only required for cartwheel assembly to regulate centriole number, but possibly it is also required to make vertical connections with the cartwheel stacks to regulate the centriole size. Specific mutations separating these two functions of CPAP would be a powerful tool to investigate mechanisms dictating this cross-talk between the centriole size and length. Recently another microcephaly mutation has been mapped in the G-box of CPAP, which resulted in missense mutation D1196N (Makhdoom *et al*, 2021). However, it is unknown how this would affect CPAP function and lead to microcephaly.

Here we have utilized two different CPAP MCPH mutations in the G-box domain, namely E1235V, and D1196N, to investigate the role of CPAP in regulating centriole length and number. Despite affecting the same G-box domain of CPAP, they caused distinct centriole phenotypes. The recently identified D1196N mutation mainly caused an increase in centriole number, and the E1235V mutation mainly caused an increase in centriole length. Interestingly, the novel D1196N mutant does not interact with the cartwheel protein STIL but could localizes to the centrioles, which indicates only a partial role of STIL in the recruitment of CPAP. Our further work points towards the involvement of a STIL-independent but CEP152-dependent alternate route, which is responsible for CPAP recruitment at the centriole. We also show that the CPAP C-terminus region (895-1338 amino acids) is sufficient for centriole localization and regulating distinct CPAP functions. Finally, we propose that such defects in centriole organization translate to multipolar spindles. Since proper spindle organization decides the division axis in neural stem cells of the neonatal brain, our work provides mechanistic insights on CPAP MCPH missense mutations leading to primary microcephaly.

## RESULTS

### CPAP is required for centriole organization

CPAP is a core centriole protein linking the proteinaceous cartwheel to the outer centriole microtubule triplet (Hatzopoulos *et al*, 2013). To investigate the effect of CPAP on centriole organization, we knocked down endogenous CPAP using the small interfering siRNA **(figure S1a and b)** and counted centriole numbers using centrin-1 marker (Paoletti *et al*, 1996). In control HeLa cells expressing EGFP fluorescent protein, the majority (84%) of interphase cells have two centrioles, and only a minor population have less than two (10 %) or more than two (6%) centrioles **(Figures 1a and b)**. However, when we depleted endogenous CPAP using siRNA, there was a significant increase (49%) in the population of interphase cells with less than two centrioles. Importantly, we could rescue this effect on centriole number by expressing ectopic siRNA-resistant EGFP tagged CPAP in the CPAP siRNA background, where 74% of interphase cells again show two centrioles. We also observed that overexpression of EGFP_CPAP does not affect the number of centrioles in interphase cells when compared to the control cells. The data suggest that CPAP is required for cartwheel assembly, thus affecting centriole numbers.

**Figure 1.**
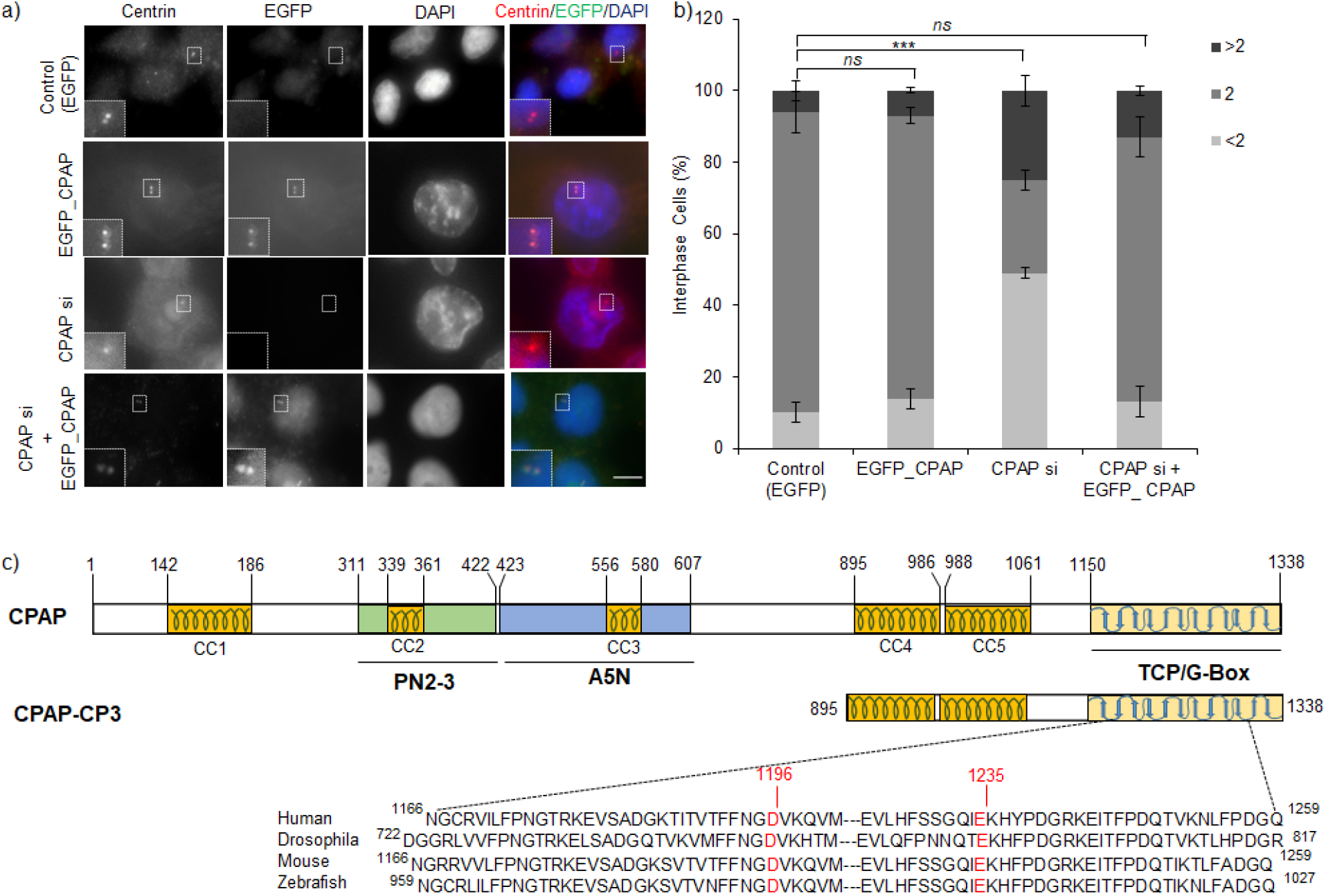
CPAP is required for centriole organization: (a) Immunofluorescence image of representative transfected HeLa cells stained for centrin-1 (centriole marker) and DAPI (nuclear marker). EGFP channel represents respective constructs in each condition. The small insert show magnified view of the centriole in respective channels. The scale bar is 5 μm. (b) Bar graph represents the percentage of interphase cells with <2 (light grey), 2 (medium grey) and >2 (dark grey) centrin-1 dots in respective condition in experiment (a). Values are average percentages ± SD from two independent experiments (n =50-100). The statistical significance represents p-value of a Chi-square test. [p>0.05: not significant (ns); p<0.001: 3 stars]. (c). The schematic representation of major domains in the human CPAP protein [PN2-3, A5N, TCP/G-box domain and five CC (coiled coil) regions]. The numbers represent amino acids marking the position of different regions in the CPAP protein. The CPAP-CP3 refers to the C-terminal (895-1338) part of CPAP comprising of CC4-5 and the G-box domain. The alignment below shows the conserved nature of MCPH-linked D1196 and E1235 amino acid residues (marked in red) in human, *Drosophila*, mouse and zebrafish.

Human CPAP is a 1338 amino acid-containing protein with three major conserved domains PN2-3, A5N, and TCP/G-box domain **(Figure 1c)**. Although the G-box domain does not interact directly with tubulin, previously reported primary microcephaly (MCPH6) missense mutations E1235V in the G-box is known to increase centriole length (MCPH6) (Bond *et al*, 2005; Kitagawa *et al*, 2011a) *via* an unknown mechanism. Recently, another primary microcephaly mutation in the G-box domain has been mapped, which substitutes aspartic acid at 1196 amino acid position with the uncharged asparagine (Makhdoom *et al*, 2021). However, the effect of D1196N mutation on CPAP function remains to be tested. We compared the amino acid sequences encompassing these two residues from organisms like humans, mice, flies, and zebrafish **(Figure 1c)**, which showed their conserved nature during evolution.

### CPAP microcephaly missense mutations in the G-box reveal separate CPAP functions in regulating centriole number and size

CPAP overexpression is known to affect centriole length with minimal effects on their numbers (Kohlmaier *et al*, 2009). We investigated the effect of CPAP microcephaly mutations on centriole number and length using centrin-1 as a centriole (number) marker and acetylated tubulin staining the centriole length in the fluorescence microscopy. HeLa cells depleted for endogenous CPAP by siRNA and expressing the RNAi-resistant wild-type EGFP_CPAP, the majority (82.5%) of interphase cells has two centrioles **(Figures 2a-b)**. Expressing EGFP_CPAP E1235V had no significant effect on the number of centrioles in the interphase cells compared to control condition. However, EGFP_CPAP D1196N expression caused more than 8-fold increase in the population of interphase cells with more than two centrioles. Since E1235V microcephaly mutation is known to affect centriole length (Kitagawa *et al*, 2011a), we investigated the effect of exogenous siRNA-resistant CPAP expression on centriole length using acetylated-tubulin marker, when the endogenous CPAP was depleted by siRNA. In accordance to previos work, we could show that E1235V expression causes an increase in centriole length **(Figures 2c-d.)**. However, D1196N had no significant effect on centriole length. The expression of CPAP-CP3 wild type and mutants in endogenous CPAP depleted state also showed the same result **(Figure S2a-d)**. This suggest that the two separate microcephaly mutations are affecting distinct functions of CPAP in regulating organelle number and length.

**Figure 2.**
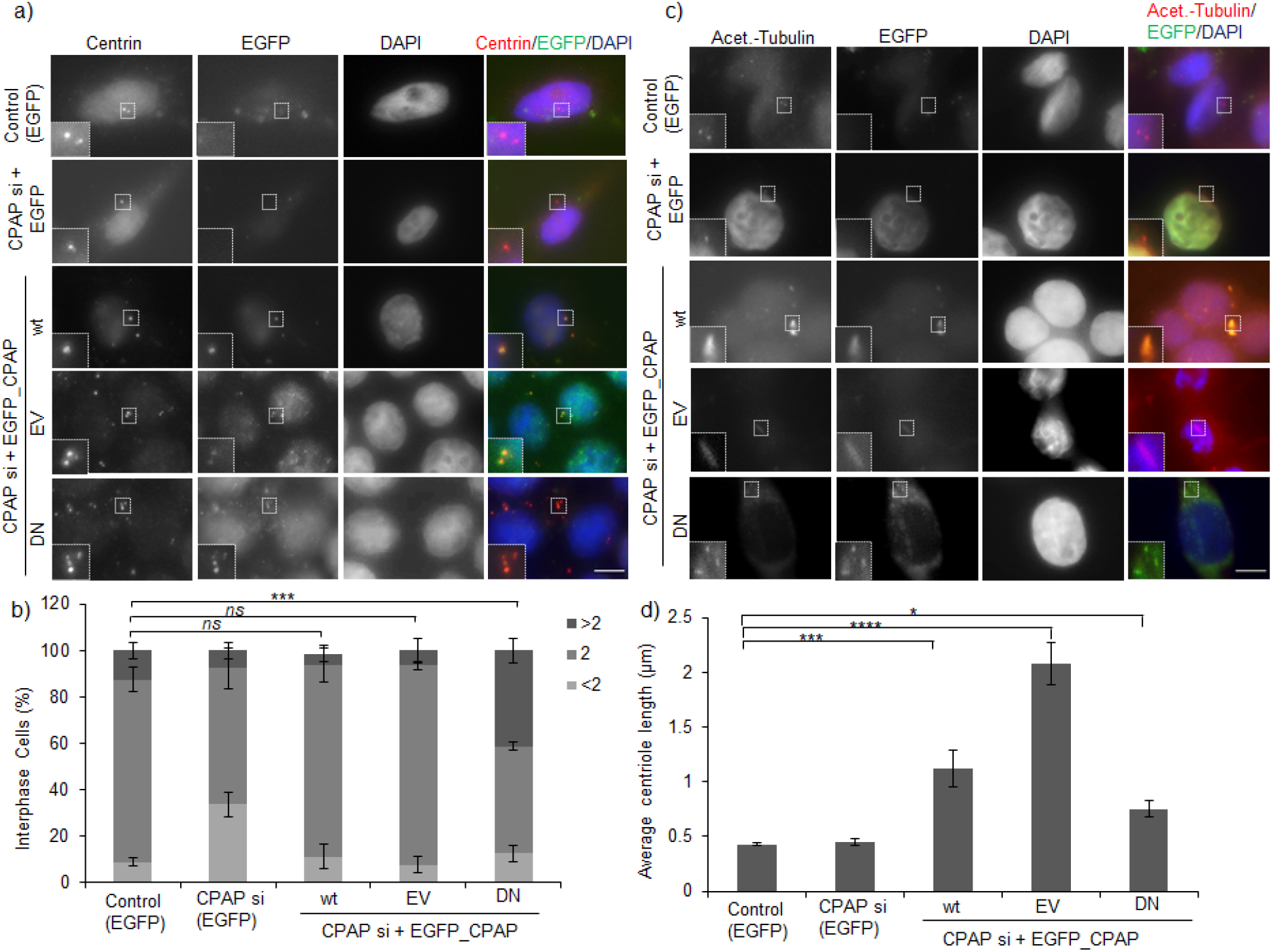
Separation of function phenotypes by CPAP MCPH, E1235V (EV) and D1196N (DN) mutations. (a) Immunofluorescence image of representative transfected HeLa cells stained for centrin-1 (centriole marker) and DAPI (nuclear marker). EGFP channel represents respective constructs in each condition. The small insert show magnified view of the centriole in respective channels. The scale bar is 5 μm. (b) Bar graphs respresenting the percentage of interphase cells with <2 (light grey), 2 (medium grey) and >2 (dark grey) centrin-1 dots in respective condition in experiment (a). Values are average percentages ± SD from two independent experiments (n =50-100). The statistical significance represents p-value of a Chisquare test. [p>0.05: not significant (ns); p<0.001: 3 stars]. (c) Immunofluorescence image of representative transfected HeLa cells stained for acetylated tubulin (centriole length marker) and DAPI (nuclear marker). EGFP channel represents respective constructs in each condition. The small insert show magnified view of the centriole in respective channels. The scale bar is 5 μm. (d) Bar graphs respresenting the average centriole length observed for the HeLa transfected with respective constructs in experiment (c). Values are average centriole length ± SD from two independent experiments (n =50-100). The statistical significance represents p-value of a student t-test. [p<0.001: 3 stars; p<0.0001: 4 stars; p<0.05: 1 star].

### CPAP E1235V (MCPH6) and novel CPAP D1196N mutations affect direct interaction with the STIL protein but vary in their centriole localization pattern

The E1235V mutation is known to disrupt CPAP interaction with the STIL protein in the *in vitro* pull-down experiment (Kitagawa *et al*, 2011a; Tang *et al*, 2009). However, the effect of novel D1196N microcephaly mutation on CPAP-STIL interaction is not known. The region of CPAP from 895-1338 amino acid residues (CPAP-CP3) is sufficient for interaction with the STIL protein in the region 375-510 amino acid residues (STIL-CR2) (Tang *et al*, 2011). Since expression of the full-length recombinant CPAP and STIL was challenging in bacteria, we used recombinant His-MBP tagged CPAP-CP3 (895-1338 amino acids) and GST-tagged STIL-CR2 (375-510 amino acids) proteins expressed in *Escherichia coli* to test their direct interaction using the GST pull-down assay. We found that just like E1235V mutation of CPAP, the D1196N mutation also significantly reduce CPAP-CP3 interaction with the STIL-CR2 region **(Figure 3a)**.

**Figure 3.**
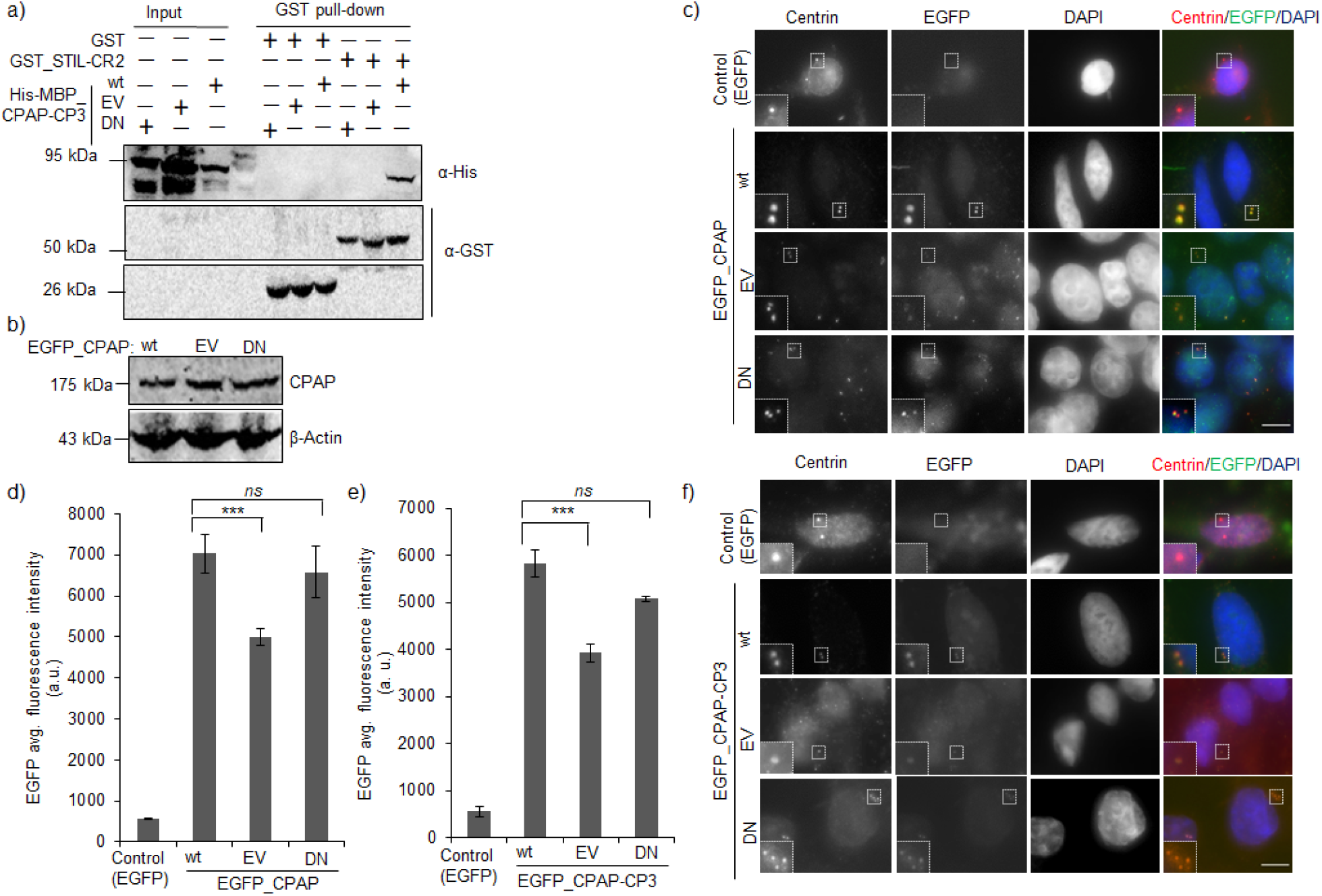
CPAP MCPH mutations, E1235V (EV) and D1196N (DN), affect direct interaction with the cartwheel STIL protein and have distinct centriole localization. (a) The western blot showing loss of interaction between GST_STIL-CR2 region and His-MBP_CPAP-CP3 E1235V (EV) and His-MBP_CPAP-CP3 D1196N (DN) as compared to the His-MBP_CPAP-CP3 wild type (wt) in the GST-pull down lanes. (b) The western blot analysis showing equal expression levels in the total cell lysates of transfected HeLa cells with EGFP_CPAP wt, EGFP_CPAP E1235V (EV) and EGFP_CPAP D1196N (DN).(c and f) Immunofluorescence image of representative transfected HeLa cells stained for centrin-1 (centriole marker) and DAPI (nuclear marker). EGFP channel represents respective constructs in each condition. The small insert show magnified view of the centriole in respective channels. The scale bar is 5 μm. (d and e) Bar graphs respresenting the average EGFP fluorescence intensity at centrioles in respective conditions of experiment (c) and (f), respectively. Values are average fluoresnce intensity ± SD from two independent experiments (n =50-100). The statistical significance represents p-value of a student t-test. [p>0.05: not significant (ns); p<0.001: 3 stars].

Next, we investigated the localization of CPAP microcephaly mutants at centrioles using fluorescence microscopy. Although, the expression of EGFP_CPAP wild type, E1235V and D1196N was comparable in transfected HeLa cells **(Figure 3b)**, we observed that the centriole localization of EGFP tagged CPAP E1235V is significantly reduced in the transfected HeLa cells as compared to the wild-type EGFP_CPAP **(Figure 3c-d.)** However, EGFP_CPAP D1196N localization was comparable to the wild-type CPAP. Similar effects were observed where only the CPAP-CP3 region (wild type or mutants) was transiently transfected in HeLa cells **(Figure 3e-f)**. This motivated us to investigate the mechanism involved in CPAP recruitment at the centrioles.

### The novel CPAP D1196N microcephaly mutation revealed an STIL-independent mechanism downstream of CEP152 involved in CPAP centriole recruitment

The STIL protein is required in CPAP localization at the procentriole (Tang *et al*, 2011). So, mutations affecting this interaction should affect the localization of CPAP, which is observed for the E1235V mutation **(Figure 3c panel 3)**. However, the novel D1196N microcephaly mutation in CPAP, which is not interacting with STIL **(Figure 2a)** can localize normally to the centrioles, which challenges the previous notion **(Figure 3c panel 4)**. To test this carefully, we depleted endogenous STIL using 90 nM siRNA **(figure S3a-b)** and observed exogenous siRNA-resistant wild type and mutant CPAP localization at centrioles by fluorescence microscopy. Depleting STIL had partial effects on the localization of exogenously transfected wild-type EGFP_CPAP at the centrioles **(Figures 4a-b.)** Similar results were obtained for the EGFP_CPAP-CP3 region **(Figures S4a-b)**. The data suggests involvement of some alternate route to STIL responsible for CPAP localization at the centrioles.

**Figure 4.**
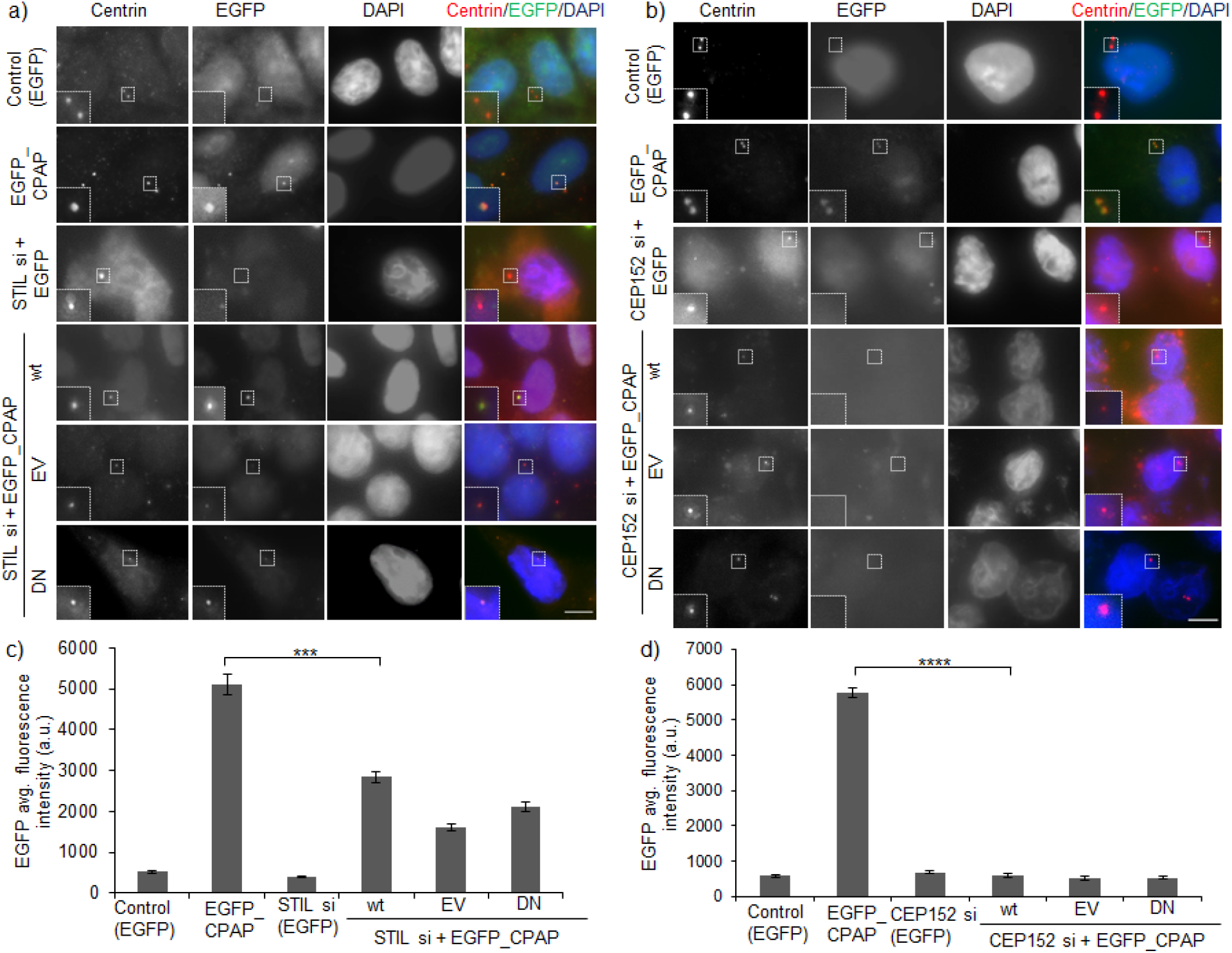
CPAP uses two alternate CEP152-dependent routes for the centriole localization. (a and b) Immunofluorescence image of representative transfected HeLa cells stained for centrin-1 (centriole marker) and DAPI (nuclear marker). EGFP channel represents respective constructs in each condition. The small insert show magnified view of the centriole in respective channels. The scale bar is 5 μm. (c and d) Bar graphs respresenting the average EGFP fluorescence intensity at centrioles in respective conditions of experiment (a) and (b), respectively. Values are average fluoresnce intensity ± SD from two independent experiments (n= 50-100). The statistical significance represents p-value of a student t- test. [p<0.001: 3 stars; p<0.0001: 4 stars].

The parent centriole protein, CEP152, is an upstream regulator shown to form a complex with the N-terminus of CPAP (Cizmecioglu *et al*, 2010). We depleted endogenous CEP152 **(Figures S3c-d)**, and observed drastict deplenishment in levels of exogenous RNAi-resistant EGFP_CPAP at the centrioles **(Figure 4c-d)**. Similar effects were observed when CEP152 was depleted in EGFP_CPAP-CP3 (wild type and mutants) expressing HeLa cells **(figure S4c-d)**. The data suggest that STIL is not working alone to recruit CPAP at centrioles, and there must be an alternate route that is CEP152-dependent. In the case of the D1196N mutant, which loses interaction with STIL protein but could normally localize to centrioles, possibly because this mutation might be enhancing recruitment through the alternate route, which remains to be tested.

### CPAP microcephaly mutations affect spindle organization in dividing cells

Proper centriole organization is essential for bipolar spindle organization and division of neural progenitor cells in developing brains. We investigated the effect of CPAP microcephaly mutants on spindle formation. We observed that the depletion of endogenous CPAP by siRNA results in a 7-fold increase in mitotic cell population with monopolar spindles (stained with α-tubulin), which supports the fact that CPAP depletion causes a decrease in centriole number **((Figures 5a-b)**. This phenotype is rescued by expressing exogenous siRNA-resistant EGFP_CPAP. However, EGFP_CPAP E1235V mutation, which causes an increase in centriole length, and D1196N mutation, which causes an increase in centriole number, showed a 3-fold increase in the population of mitotic cells with the multipolar spindle. This suggests that although the two microcephaly mutations in the G-box of CPAP control two aspects of its functioning, they ultimately cause similar spindle organization problems, which could explain their contribution to the etiology of primary microcephaly by affecting neural stem cell division in the neonatal brain.

**Figure 5.**
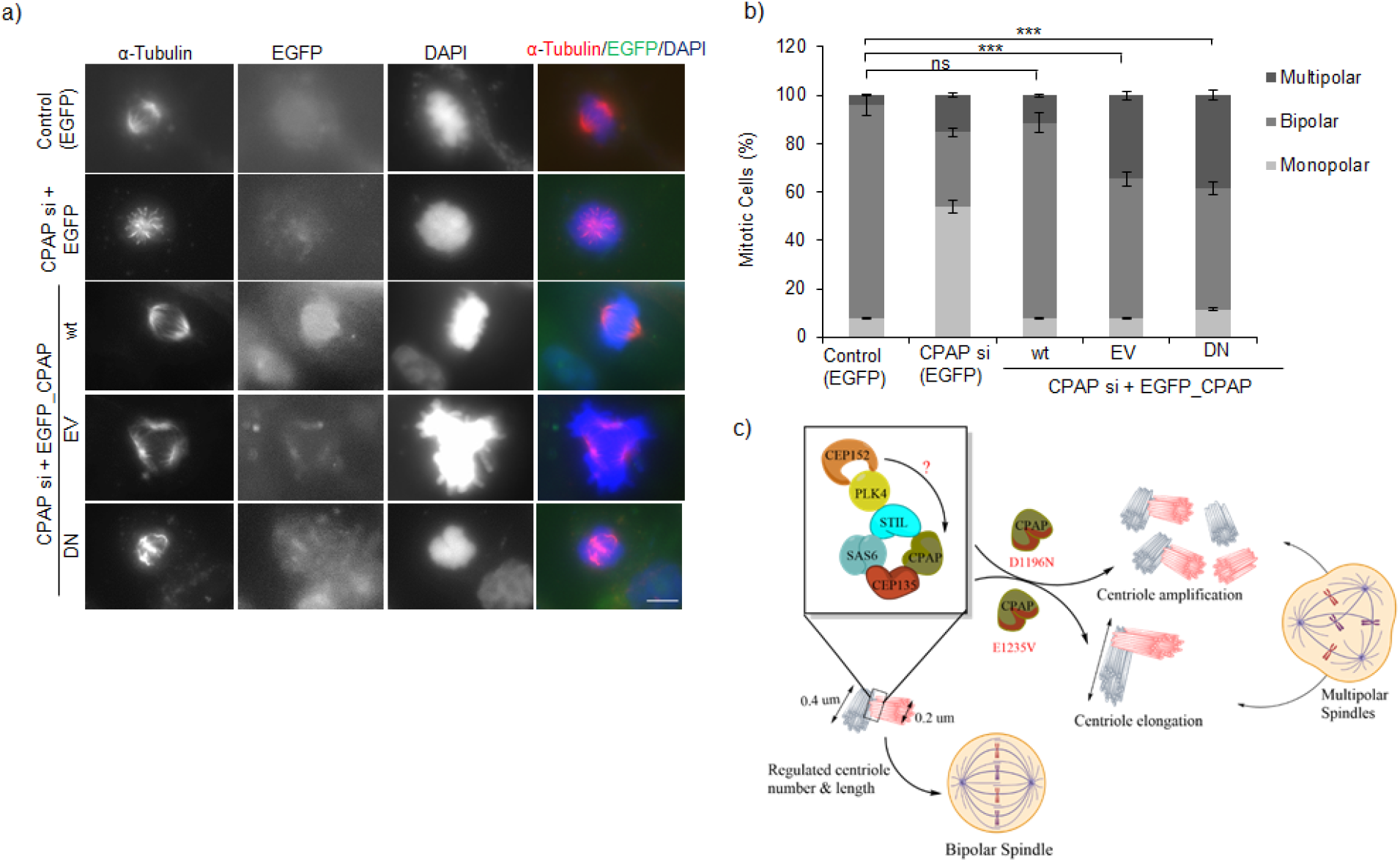
CPAP MCPH mutation causes an increase in mutilpolar spindles in mitotic cells. (a) Immunofluorescence image of HeLa cells transfected with respective construts and synchronized at M phase to visualize mitotic spindle using α-tubulin as marker and and DAPI (nuclear marker). EGFP channel represents respective constructs in each condition. The scale bar is 5 μm. (b) Bar graphs respresenting the percentage of mitotic cells with monopolar (light grey), bipolar (medium grey) and multipolar (dark grey) spindles in respective conditions quantified from experiment (a). Values are average percentages ± SD from two independent experiments (n=50). The statistical significance represents p-value of a student t-test. [p>0.05: not significant (ns); p<0.001: 3 stars]. (c) The proposed model, showing unknown CEP152-dependent alternate route utilized by CPAP for centriole recruitment. The two MCPH mutations in CPAP cause distinct effects on CPAP functioning resulting in abnormal spindle organization in a dividing cell.

## DISCUSSION

The core structure of centrosome are made of microtubule-based structures called centriole. The centriole microtubules are more stable and slow growing than the cytoplasmic microtubules. However, it remains puzzling as to how centrioles achieve definite size and stability. CPAP has emerged as an essential protein linking central cartwheel proteins to the growing outer microtubules and possibly also stacking these cartwheels along the length of the centriole lumen. The CPAP microtubule destabilizing, PN2-3, and stabilizing A5N domains are known to interact directly with centriole microtubules to regulate their growth and length. The c-terminal G-box domain of CPAP is engaged in direct interaction with the core centriole cartwheel protein STIL, which is known to be responsible for the recruitment of CPAP and procentiole generation. Two separate primary microcephaly missense mutations have been mapped in the G-box of CPAP, thus suggesting the importance of this domain in the functioning of CPAP. Although the G-box domain of CPAP falls outside the microtubule/tubulin binding region, we show that it is necessary for controlling distinct CPAP functions.

Here we simultaneously characterized the two reported MCPH mutations in the G-box of CPAP (E1235V, and D1196N), which revealed that despite affecting the same domain of the proteins, they have different effects on CPAP function. According to previous literature, the CPAP E1235V (MCPH6) overexpression caused overly long centrioles with no effect on the centriole number. Interestingly, characterization of the recently identified CPAP D1196N microcephaly mutation revealed that its overexpression does not affect centriole length. However, it causes a significant increase in the centriole number. We show that both the CPAP, E1235V and D1196N mutations affect the CPAP direct interaction with the STIL protein in the *in-vitro* pull-down experiment. Moreover, we observed using fluorescence microcopy that E1235V mutation affects CPAP localization at the centrosome. However, D1196N mutant localization remains unaffected when compared to the wild-type CPAP. Our work revealed that STIL is only partially required for the localization of CPAP at the centrosome, and there is an alternative CEP152-dependent pathway that is simultaneously needed for the recruitment of CPAP to the centrosomes **(Figure 5c)**. This work highlights the importance of the CPAP G-box domain in regulating two distinct functions of CPAP, i.e., centriole number and centriole size/length. Since important functional mutations map to the G-box domain of CPAP, further work is required to understand the involvement of this region in CPAP activity. In future, it would be interesting to investigate and compare the structural effect of CPAP E1235V and D1196N microcephaly mutation on the G-box organization in relation to tubulin. In addition, both investigated MCPH mutations substitute negatively charged amino acids for uncharged amino acids, which points towards the involvement of protein interactions. Several centriolar proteins found in complexes with CPAP, like, CEP135 (Lin *et al*, 2013a), Centrobin (Gudi *et al*, 2011), and CEP120 (Lin *et al*, 2013b), are also involved in regulating centriole length. These would be attractive candidates for gaining mechanistic insights and their possible involvement in the CPAP centriole localization and functioning.

## MATERIAL & METHODS

### Molecular Biology

The siRNA-resistant CPAP cDNA containing plasmid was obtained from Addgene (#46390). Full-length and C-terminal coding sequences (residues 895-1338 CP3) were PCR amplified from the siRNA-resistant CPAP, which were fused in the same reading frame to EGFP in pCDNA5/FRT/TO or to 6xHis-MBP tag in pET Duet vector. Full-length and mutants (E1235V and D1196N) were generated by site-directed mutagenesis using the Phusion High-Fidelity DNA Polymerase (New England Biolabs, #M0530S), followed by DpnI digestion (Takara, #1235A). The cDNA of STIL ((Addgene, #80266) and the CR2 fragment (residues 375-510) sequences was PCR amplified and cloned in-frame to the GST tag in pGEX-6P1 vector.

### Protein expression and purification

E. coli (BL21) cells were transformed with the recombinant plasmid pETDuet_His-MBP_CPAP-CP3 (wt and mutants) and pGEX6P1_GST_STIL-CR2. The bacterial cultures were grown at 37 °C until an OD_600_ of 0.4-0.8. Protein expression was induced by adding either 0.4 mM IPTG to the cell culture with pET_Duet_His-MBP_CPAP-CP3 (wt and mutants) or 1 mM IPTG, 0.1 mM ZnCl_2_, 2% Glucose solution and 1.5% Ethanol solution for pGEX6P1_GST_STIL-CR2. The cells were grown at 18°C for 18 hours. For affinity purification of the His-tagged proteins, the bacterial cells were lysed using the lysis buffer (50 mM Tris-HCl pH 8.0, 300 mM NaCl, 5 mM Imidazole). The cell lysate was sonicated at 35% amp on ice thrice for 10 sec with 10 sec break. Th cell debris was pelleted down at 13000 rpm, 10 minutes 4°C and the supernatant was incubated with pre-equilibrated Ni-NTA agarose beads (Qiagen, #1018244) with shaking on ice for 3 hours. The beads were spun down at 2000 rpm for 2 min to remove unbound fraction. This was followed by three washing steps with the washing buffer (50 mM Tris-HCl pH 8.0, 300 mM NaCl, 20 mM Imidazole). Finally, the protein was eluted from the beads using the elution buffer (50 mM Tris-Cl pH 8.0, 300 mM NaCl, 300 mM Imidazole). GST tagged proteins were purified by the same protocol using the GST-lysis buffer (50 mM Tris-HCl pH 8.0, 100 mM NaCl, 1% Triton-X, 1mM DTT, 1 mM PMSF), the gluthathione superflow resin (Takara, #635607), and the GST-wash buffer (1x PBST, 1 % NP-40, 2 mM DTT).

### Cell Line, Transfection, and Media

HeLa Kyoto cells were cultured in DMEM medium (Himedia,#AL111) supplemented with 10% fetal bovine serum, (Merck,#12103C), 2 mM antibiotics (penicillin and streptomycin) solution (Merck, #P4333), and 2 mM L-glutamine (Merck, #G7513) in a humidified incubator with 5% CO_2_ at 37 °C. GFP-tagged cDNA constructs and siRNAs were transiently transfected using the Lipofectamine 3000 (Invitrogen, #L3000001) as per manufacturer’s protocol. Double stranded CPAP siRNA oligonucleotides was synthesized with 3’-UU overhangs with the sequence 5’- AGAAUUAGCUCGAAUAGAA-3 ‘ (Dharmacon). STIL and CEP152 siRNA duplexes were synthesized with the sequence 5’- GUUUAAGGGAAAAGUUAUU-rU-rU-3 ‘ and 5’- GGAGGACCAUGUCAUUAGACU-rU-rU-3’, respectively (Merck).

### Antibodies and siRNAs

Primary antibodies used for immunoblotting were rabbit anti-CenpJ (Invitrogen,#PA597577, 1:1000), mouse anti-His (Invitrogen, #MA1-21315, 1:1000), rabbit anti-GST (Merck,#ZRB1223,1:1000), mouse anti-β-actin (Santa Cruz Biotechnology,#SC-47778,1:1000). Secondary antibodies used were anti-mouse IgG HRP linked (Cell Signaling Technology,#CST 7076S,1:10,000) and anti-Rabbit IgG HRP linked (Cell Signaling Technology, CST 7074S, 1:10,000). Primary antibodies used for immunofluorescence were mouse anti-centrin1 (Merck, #04-1624, 1:500), rabbit anti-STIL (Abcam, #Ab89314, 1:100), mouse anti-α tubulin (Merck, #T6074, 1:500), mouse anti-tubulin acetylated (Merck, #T6793), rabbit anti-CENPJ (Invitrogen,#PA597577, 1:300). Secondary antibodies used were anti-rabbit IgG (Fc) TRITC (Merck, # SAB3700846, 1:500) and anti-mouse IgG (Fc) TRITC (Merck, #SAB3701020, 1:500).

### Transfection, and Drug Treatment

For plasmid transfections, cells at 80–90% confluency were transfected using Lipofectamine 3000 as per manufacturer’s protocol and analyzed after 48h. To visualize metaphase cells, they were synchronized for G_2_/M phase by using 5 uM RO-3306 (Merck, #SML0569) for 18h followed by 45 minutes of release. Cells were synchronized at the S-phase using the double-thymidine block (2 mM) to detect endogenous STIL. Cells were incubated with thymidine for 18 hours followed by 9 hours of release and the second block was introduced for additional 15 hours followed by cell fixation for immunofluorescence.

### Immunofluorescence

HeLa cells grown on coverslips were washed with 1X PBS to remove media. They were permealized using 0.1% TritonX-100 prepared in 1XPBS (1XPBST) and fixed with 4% (v/v) paraformaldehyde solution prepared in 1X PBS for 10 min at room temperature. The cells were washed thrice with 1XPBST and blocked using 1% BSA in 1XPBST for 1 h. The cells were washed thrice with 1XPBST and left in primary antibodies overnight at 4 °C. After primary antibody incubation, cells were washed thrice and incubated in secondary antibody for an hour at room temperature. Cell were stained with DAPI (0.5 ug/ml) prepared in 1x PBS for 15 minutes and mounted on glass slides with the help of mouting solution (0.62 g DABCO, 2 g mowiol, 6.25 ml glycerol, 25 ml 0.2 M Tris-Cl pH 8.5). The fluorescent images were acquired using the 100X oil, 1.35 NA objective lens of the epi-fluorescence microscope (Olympus IX83). Images are represented as maximum projections of Z-stacks and analyzed using ImageJ. The bar graphs were ploted using the Microsoft excel software and figures were arranged on Microsoft powerpoint.

### GST-pull down assay

GST pull-down assay was performed to test the interaction between 6xHis-MBP tagged CPAP CP3 wild-type/mutant proteins with the GST_STIL-CR2 region. Bacterial cell lysates expressing His-MBP_CPAP-CP3 constructs were incubated with immobilized GST_STIL-CR2 fusion protein at 4 °C for 3 h in binding buffer (50 mM Tris-HCl pH 8.0, 100 mM NaCl, 1% Triton-X, 1mM DTT, 1 mM PMSF). After incubation, the beds were washed three times with the GST-wash buffer. The proteins were eluted from the beads by incubating with the SDS lamelli buffer and followed by boiling at 95 °C for 10 min. The proteins were resolved on a 10% SDS–PAGE. The protein bands were transferred to nitrocellulose membrane and analyzed by western blotting (Sambrook, 2001).

## ACKNOWLEDGEMENT

The HeLa Kyoto, pCDNA5 /FRT/TO, pETDuet1 and pGEX vectors were kind gift from the laboratory of Prof. Andrea Musacchio, Max Planck Institute of Molecular Physiology, Dortmund, Germany. We are thankful to Dr. Stefano Maffini and Dr. Clemens Cabernard for critical reading of the manuscript. SJ and SS are supported by GATE fellowship from the Ministrity of Education, Government of India. The work is supported by the grants received from Science and Engineering Research Board (ECR/2017/001410), Department of Biotechnology (BT/12/IYBA/2019/02) and Board of Research in Nuclear Sciences (55/14/02/2021-BRNS/10206)

## AUTHORS CONTRIBUTION

SJ and PS conceptualized the work. SJ performed major experiments with the help of SS. All authors contributed to the data analysis and writing of manuscript.

