## Supplementary Data for "Separation of function mutations in microcephaly protein CPAP/CENPJ, reveal its role in regulating centriole number and length"

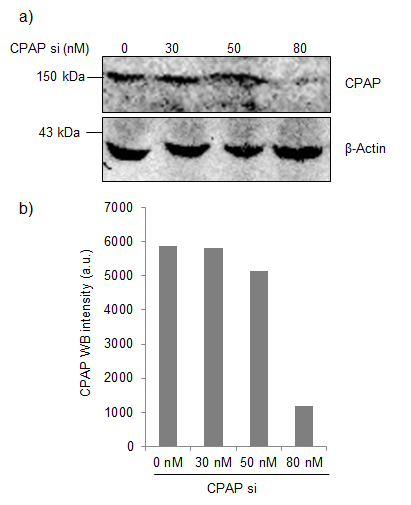

**Figure S1**. Optimization of endogenous CPAP knockdown in HeLa cells. (a) Western blot analysis of total cell lysate showing knockdown of endogenous CPAP using siRNA. Significant knockdown was observed at 90 nM, which was used for subsequent experiments. (b) Quantification of CPAP western blot band intensity in the experiment (a).

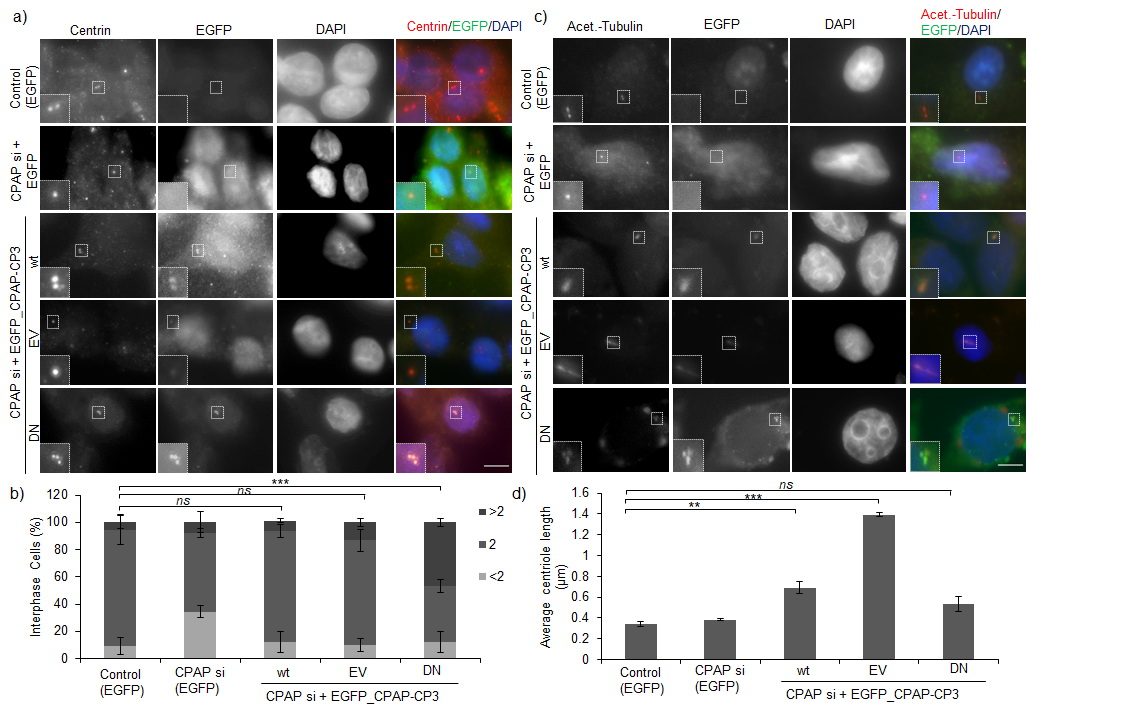

**Figure S2**. MCPH mutations in the C-terminus of CPAP affect distinct functions in regulating centriole number and length. (a) Immunofluorescence image of representative transfected HeLa cells stained for centrin-1 (centriole marker) and DAPI (nuclear marker). EGFP channel represents respective constructs in each condition. The small insert shows magnified view of the centriole in respective channels. The scale bar is 5 μm. (b) Bar graphs representing the percentage of interphase cells with <2 (light grey), 2 (medium grey) and >2 (dark grey) centrin-1 dots in the respective conditions of the experiment (a). Values are average percentages ± SD from two independent experiments (n>50). The statistical significance represents p-value of a Chi-square test. [p>0.05: not significant (ns); p<0.001: 3 stars]. (c) Immunofluorescence image of representative transfected HeLa cells stained for acetylated-tubulin (centriole length marker) and DAPI (nuclear marker). EGFP channel represents respective constructs in each condition. The small insert shows magnified view of the centriole in respective channels. The scale bar is 5 μm. (d) Bar graphs representing average centriole length observed for the HeLa transfected with the respective constructs in the experiment (c) Values are average centriole length ± SD from two independent experiments (n>50). The statistical significance represents p-value of a student t-test. [p>0.05: not significant (ns); p<0.01: 2 stars; p<0.001: 3 stars].

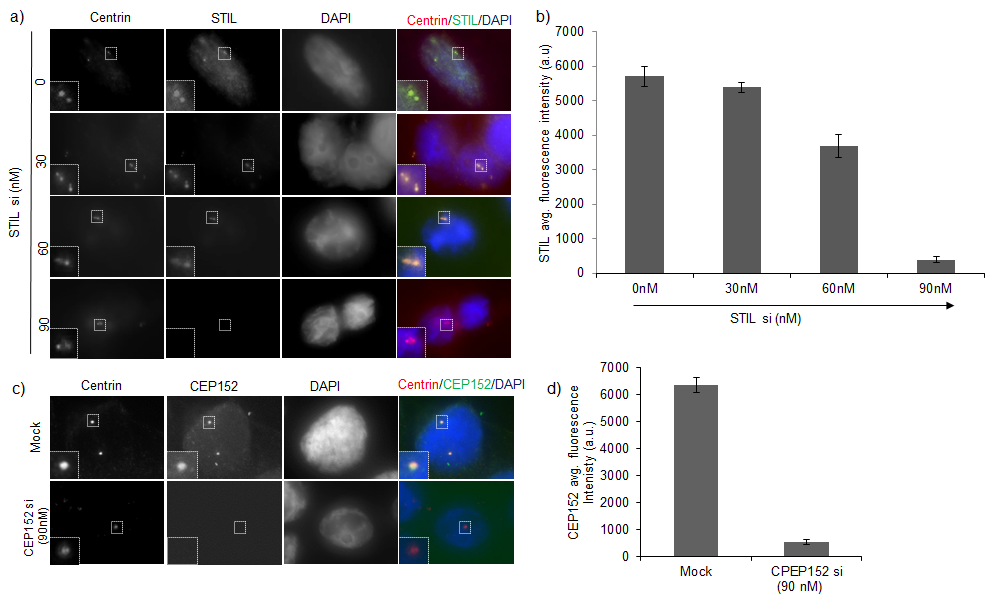

**Figure S3**. Optimization of endogenous STIL and CEP152 knockdown in HeLa cells. (a) Immunofluorescence image showing dose-dependent decrease in the STIL protein levels at the centrioles in the respective siRNA treated HeLa cells. Small inserts provide magnified view of the centrioles. Significant reduction in STIL level at the centrioles was observed at 90 nM, which is used for the subsequent experiments. The scale bar in 5 μm. (b) Quantification of the STIL protein absolute fluorescence intensity at centrioles. (c) Immunofluorescence image showing reduction in CEP152 protein at centrioles on knockdown with 90 nM siRNA. Small inserts provide magnified view of the centrioles. The scale bar in 5 μm. (b) Quantification of the CEP152 protein absolute fluorescence intensity at the centrioles.

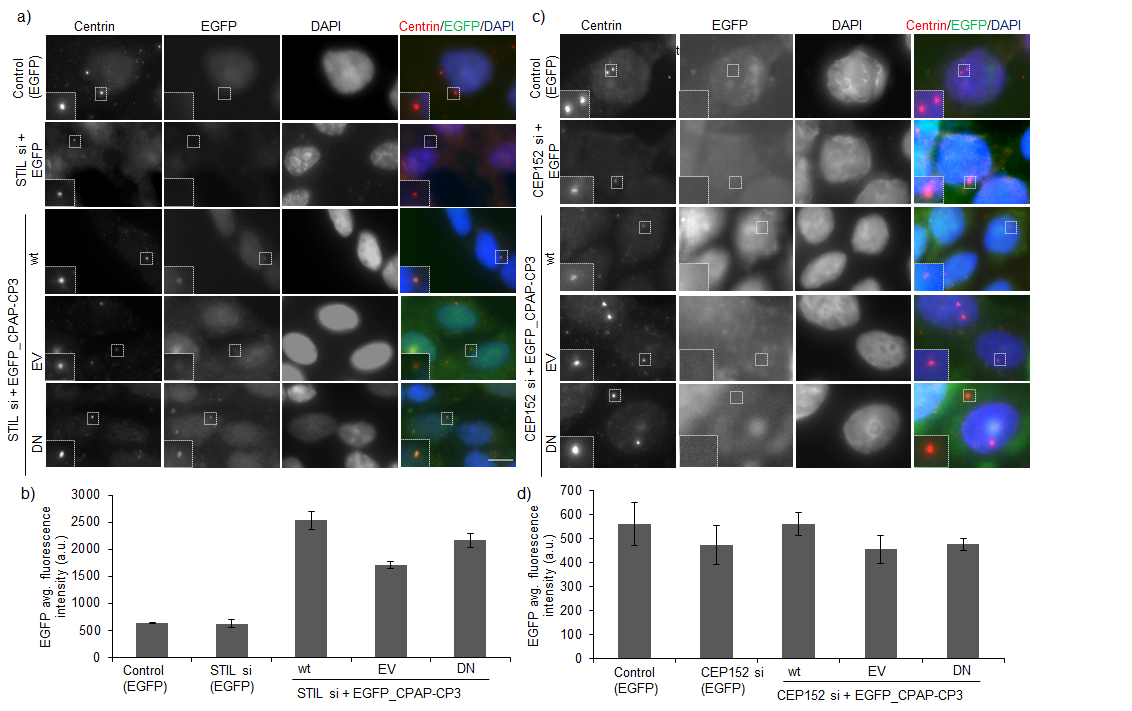

**Figure S4.** CPAP-CP3 domain localization at centriole is partially dependent on STIL and uses an alternate CEP152-dependent route for centriole localization. (a and c) Immunofluorescence image of representative transfected HeLa cells stained for centrin-1 (centriole marker) and DAPI (nuclear marker). EGFP channel represents respective constructs in each condition. The small insert shows magnified view of the centriole in the respective channels. The scale bar is 5 μm. (b and d) Bar graphs representing the average EGFP fluorescence intensity at the centrioles in the experiment (a) and (c), respectively. Values are average fluorescence intensity ± SD from two independent experiments (n>50).
